# Boosting the Sensitivity of Quantitative Single-Cell Proteomics with Activated lon-Tandem Mass Tags (AI-TMT)

**DOI:** 10.1101/2024.02.24.581874

**Authors:** Trenton M. Peters-Clarke, Yiran Liang, Keaton L. Mertz, Kenneth W. Lee, Michael S. Westphall, Joshua D. Hinkle, Graeme C. McAlister, John E. P. Syka, Ryan T. Kelly, Joshua J. Coon

**Affiliations:** Department of Chemistry, University of Wisconsin-Madison, Madison, WI, 53706, USA; Department of Biomolecular Chemistry, University of Wisconsin-Madison, Madison, WI, 53706, USA; Department of Chemistry, Brigham Young University, Provo, UT, 84602, USA; Thermo Fisher Scientific, San Jose, CA, 95134, USA; National Center for Quantitative Biology of Complex Systems, Madison, WI, 53706, USA; Morgridge Institute for Research, Madison, WI, 53515, USA

## Abstract

Single-cell proteomics is a powerful approach to precisely profile protein landscapes within individual cells toward a comprehensive understanding of proteomic functions and tissue and cellular states. The inherent challenges associated with limited starting material in single-cell analyses demands heightened analytical sensitivity. Just as advances in sample preparation maximize the amount of material that makes it from the cell to the mass spectrometer, we strive to maximize the number of ions that make it from ion source to the detector. In isobaric tagging experiments, limited reporter ion generation limits quantitative accuracy and precision. The combination of infrared photoactivation and ion parking circumvents the *m/z* dependence inherent in HCD, maximizing reporter generation and avoiding unintended degradation of TMT reporter molecules in a method we term activated ion-tandem mass tags (AI-TMT). The method was applied to single-cell human proteomes using 18-plex TMTpro, resulting in a 4-5-fold increase in reporter ion signal on average compared to conventional SPS-MS^3^ approaches. AI-TMT enables faster duty cycles, higher throughput, and increased peptide identification and quantification. Comparative experiments showcase 4-5-fold lower injection times for AI-TMT, providing superior sensitivity without compromising accuracy. In all, AI-TMT enhances the sensitivity and dynamic range of proteomic experiments and is compatible with other techniques, including gas-phase fractionation and real-time searching, promising increased gains in the study of cellular heterogeneity and disease mechanisms.

## INTRODUCTION

Single-cell proteomics (SCP) has emerged as a powerful approach to unravel the intricate proteomic landscapes of individual cells, enabling insights into cellular heterogeneity, developmental processes, and disease mechanisms.^1–4^ The ability to interrogate the proteome at the single-cell level holds tremendous potential for a more granular deciphering of complex biological systems. However, the inherent challenges associated with limited sample material in single-cell analysis demand highly sensitive analytical techniques.

While the prospect of SCP was once a distant goal,^5^ advances in instrumentation and data acquisition strategies have made SCP a reality.^3,4,6–10^ Developments in sample processing that minimize protein loss have been crucial to maximize the amount of peptide ions entering the mass spectrometer.^6,7,11–13^ Isobaric labeling approaches with tandem mass tags (TMTs) have become a centerpiece of SCP through methods like SCoPE-MS, which use carrier channels to boost MS^1^-level detection and reporter ions in MS/MS scans to provide relative quantitation of labeled proteins of individual cells.^6^ TMT enables sample multiplexing and quantification of proteomes across experimental conditions.^14–19^

Despite this strong progress, co-isolation of multiple precursor ions remains a major challenge of isobaric labeling strategies. Co-isolation of background species with target peptides results in reporter ion ratio distortion.^20^ To improve reporter ion tag purity, Q-Orbitrap-quadrupole linear ion trap (QLT) systems enable a synchronous precursor selection (SPS)-MS^3^ strategy, where peptides are first fragmented in the QLT by ion trap collisionally activated dissociation (CAD) and then multiple product ions are simultaneously selected by a multi-notch waveform for beam-type CAD (HCD) activation and Orbitrap MS^3^ mass analysis.^21^ This reduces distortion arising from co-isolation, significantly improving quantitative results.^21,22^ Because the optimal voltage for “beam-type” HCD increases linearly with precursor *m/z*, simultaneous activation of synchronously isolated products that span an m/z range often generates suboptimal reporter intensities.^23^ We have previously shown that a combination of photoactivation and ion parking maximizes reporter generation from tagged peptides by 1) eliminating the *m/z* dependence inherent in HCD and 2) avoiding unintended degradation of TMT reporter.^23^ We applied this method to a diluted yeast proteome and to single-cell human proteomes to highlight quantitative sensitivity improvements for limited sample amounts.

In single-cell proteomics, the limited starting material produces low ion counts and challenges in detecting low-abundance peptides.^1,8^ Just as advances in sample preparation maximize the amount of material that makes it from the cell to the mass spectrometer, we strive to maximize the number of peptide ions that make it from ion source to the detector.^12,13^ Herein, we report activated ion-TMT (AI-TMT), an MS^3^-based quantitative proteomics method employing infrared multiphoton photoactivation and reporter ion parking to maximize sensitivity and accuracy of isobaric tagging studies. As an extension of our previous work, we applied IR photoactivation and an RF ion parking waveform to TMTpro labeled peptides and applied the method to single-cell proteomics to greatly boost the generation of quantitative reporter ions. By optimizing TMT reporter ion generation within SPS-MS^3^ workflows, researchers can maximize the utilization of available ions and significantly enhance sensitivity. We report 4-5-fold higher reporter ion signal, on average, with AI-TMT over the conventional SPS-MS^3^ approach employing HCD. This leads to more than twice as many peptides quantified as HCD in at least one channel and over ten times as many peptides quantified as HCD across all 18 channels in single-cell proteomics experiments of human cell lines using 18-plex TMTpro.

This method provides several major advances for the field of quantitative proteomics. Enhanced sensitivity boosts the dynamic range of quantifiable peptides and improves quantitative accuracy, reducing measurement variability. Higher reporter ion intensity also enables shorter ion injection times, improving duty cycle and therefore the number of identified and quantitified peptides. Further, AI-TMT is perfectly compatible with other SCP and isobaric labeling methods, including gas-phase fractionation (e.g. FAIMS) which further reduces ion co-isolation, real-time searching (RTS) which increases MS efficiency, and increased multiplexing abilities as TMT methods expand to 18-plex and 34-plex workflows.

## EXPERIMENTAL SECTION

### Materials

Pierce TMT-11plex yeast triple-knockout standard (Thermo Fisher Scientific) was reconstituted in 0.2% formic acid to 1 μg/μL and was serial diluted for injection of 1 ng to 500 ng peptide. Acetonitrile (HPLC grade), H2O (HPLC grade), and formic acid were obtained from Sigma-Aldrich (St. Louis, MO).

Single-cell sample preparation using nanoPOTS. A nested nanoPOTS (N2) glass chip with 16 nanowells nested in per well was fabricated using standard photolithography and wet chemical etching as described previously.^24–26^ A regular microscope glass slide was glued with a 1-mm-thick PMMA frame to serve as the cover for long-term incubation.

HeLa and A549 cells (ATCC, Manassas, VA) were cultured at 37 °C with 5% CO2 using Dulbecco’s Modified Eagle Medium (DMEM) cell culture medium with 10% fetal bovine serum and 1% penicillin/streptomycin and were harvested upon 70% confluency. Before collection, the cells were rinsed 3 times with iced phosphate buffered saline (PBS).

The TMTpro-126 labeled carrier samples and the TMTpro-134N labeled reference samples were prepared and labeled in bulk using S-trap as described elsewhere.^24,27^ The final concentration was determined using the Pierce quantitative peptide assay. The single cell channels were prepared separately in each nanowell using the cellenONE platform (Cellenion, Lyon, France). The temperature was set to 1 °C below the dew point in the cellenONE software, and the humidity was set to 40% before the glass chip was mounted on the cellenONE stage. Within each nested well, 14 single cells and 1 PBS droplet were sorted into each nanowell. Then, 10 nL of 5 mM TCEP (Thermo Fisher Scientific, Waltham, MA) containing 0.05% n-dodecyl-β-d-maltoside (DDM) in 100 mM 4-(2-hydroxyethyl)-1-piperazineethanesulfonic acid (HEPES) at pH = 8.5 was added followed by 5 nL of 45 mM chloroacetamide. The glass chip was then clamped with its cover and incubated at 70 °C for 30 min and then at 95 °C for 15 min in a wet box to minimize evaporation. The wet box was then placed at 4 °C for 15 min. After cooling down, the chip was mounted back in the cellenONE and 5 nL of mixed solution containing 0.25 ng of Lys-c and 0.5 ng of trypsin in 100 mM HEPES (pH = 8.5) was added into each nanowell followed by the 12 h digestion at 37 °C. After incubation, the temperature of the stage was set to 20 °C and 1 nL of dimethyl sulfoxide (DMSO) was added into each nanowell followed by 5 nL of 10 g/L corresponding TMTpro reagents. The temperature was then set to 25 °C for 1 h incubation. Later, 2 nL of 5% hydroxylamine solution was dispensed into each nanowell for 15 min quenching. The samples were acidified using 5 nL of 5% formic acid before 0.5 ng of 134N-labeled HeLa and A549 digest (1:1) and 10 ng of 126-labeled HeLa and A549 digest (1:1) were added to each nested-well. At last, the samples in each nested-well were pooled and rinsed with 2.5 μL of 0.1% formic acid respectively and collected into the same well on a 384 PCR plate.

### LC-MS/MS

Experiments were performed on a quadrupole-Orbitrap-ion trap mass spectrometer modified to include a 40 W continuous-wave laser which allowed for photoactivation of ions within the ion trap.^28,29^ Adjustments to the instrument control code allowed for broadband ion parking at reporter ion m/z 126–131 during IRMPD experiments to avoid IR-induced degradation of reporter.^23^ Data-dependent MS^2^ scans were followed by standard HCD or IRMPD MS^3^ scans of SPS isolated MS^2^ product ions. MS/MS activation was performed with either HCD or CAD with scans analyzed in the Orbitrap or linear ion trap, respectively, as denoted in the text. For HCD MS/MS experiments, Orbitrap MS^1^ scans were performed at 60,000 resolving power (at *m/z* 200), MS^2^ scans were performed at 30,000 resolving power, and MS^3^ scans were performed at 60,000 resolving power. For CAD MS/MS experiments, Orbitrap MS^1^ scans were performed at 60,000 resolving power, ion trap MS^2^ scans were performed using the rapid scan rate, and Orbitrap MS^3^ scans were performed at 60,000 resolving power. Ion injection times were set to default values unless otherwise noted in the text. Columns with 50 μm I.D. were packed at 30,000 psi.^30^ A flow rate of 140 nL/min was used during liquid chromatography (LC) separations. Figure 1 summarizes the single-cell proteomics preparation and AI-TMT workflow and benefits for abundant or non-abundant peptides.

**Figure 1.**
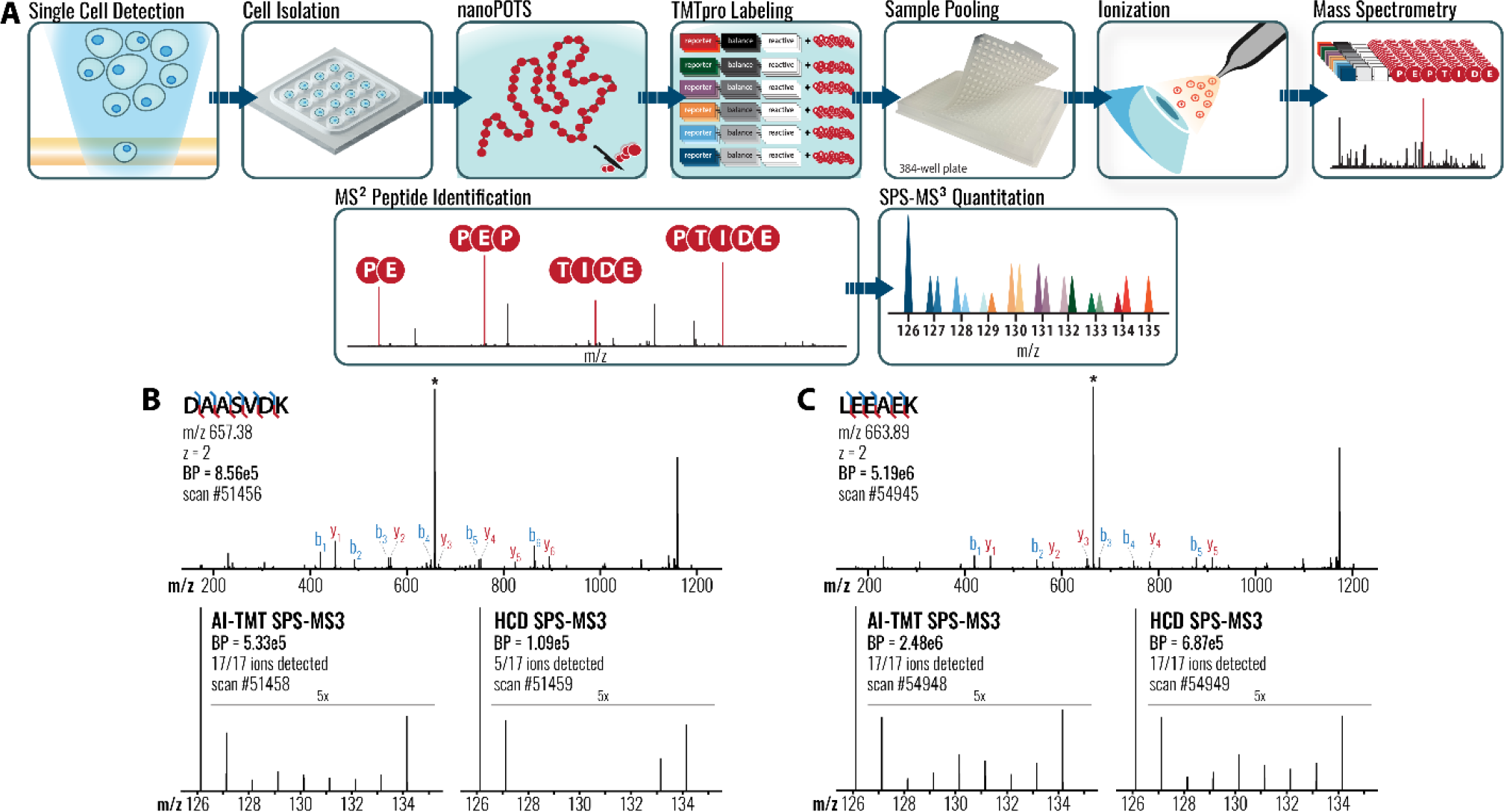
Single-cell proteomics workflow with MS^3^-based AI-TMT quantification. **A**, Illustration of the single-cell proteomics sample preparation and data acquisition workflow, from nanoPOTS and TMTpro labeling to AI-TMT quantification. **B**, Identification of a low-abundance peptide ion and quantification with AI-TMT or HCD SPS MS^3^. **C**, Identification of a high-abundance peptide ion and quantification with AI-TMT or HCD SPS MS^3^.

### Data Analysis

Mass spectra and chromatogram information were accessed in the vendor’s post-acquisition software (Xcalibur Qual Browser, version 4.0). No microscans were performed. Peak lists and intensities from XTRACT were input into the Interactive Peptide Spectral Annotator (IPSA) for sequencing and fragmentation efficiency metrics.^31^ All fragment matches were made within a 10 ppm tolerance. A Python script was written using RawQuant^32^ to directly extract precursor, fragment, and reporter ions and their corresponding intensities (in terms of signal-to-noise). Additionally, a code was written in R to plot parameter optimization and its corresponding statistics. The OMSSA algorithm and COMPASS software suite were used for searching and processing data.^33,34^ Peptides were searched with a 10 ppm tolerance around the monoisotopic precursor and a 10 ppm tolerance on fragment ion masses. Search results were filtered to a 1% unique peptide FDR based on expectation value (E-value) and ppm error using COMPASS.^33,35^ For peptide quantification, we required a TMT reporter ion S/N above zero in at least one channel. Rather than filter based on reporter ion S/N, we report the S/N values and number of channels quantified (S/N > 0) across all channels for each experiment.

## RESULTS AND DISCUSSION

### AI-TMT at low peptide loading amounts

As reporter ions are susceptible to unintended fragmentation and scattering by high-energy collisions, we photoactivated ten b- or y-type ions and prevented successive dissociation of generated reporter ions with ion parking, which we anticipated would aid in the sensitivity of TMT-based quantitation. Reporter ion yield increased by 200%, on average, for 1.2 μg peptide loading. However, when only 10 ng yeast peptide were injected, photoactivation produced a 398% average boost for the cumulative TMT reporter intensity across all eleven channels (**Figure 2**). This illustrates the inefficiency of HCD when activating multiple peptide fragments at once^23^, a phenomenon made evident for low-abundance precursors. We suspect that this boost in AI-TMT performance relative to HCD for low abundance samples comes at the limit of detection. In the Orbitrap, we can’t detect peaks with less than ∼5-10 ions, so the low intensity values of HCD are seemingly rounded down to zero and the average boost in signal with AI-TMT vs. HCD increases. For this back-to-back experiment, photoactivation quantified 38% more peptides across all 11 channels (11,280 vs. 8,151) and there were no peptides that HCD fully quantified that photoactivation didn’t already fully quantify (**Figure 3**). The criteria for peptide quantification were reporter ion S/N values above zero across all channels. The 3,129 peptides fully quantified by only IR photoactivation are all among the lowest abundant peptides (**Figure 3d**). Our method allows researchers to quantify progressively low abundance peptides by boosting TMT reporter intensity.

**Figure 2.**
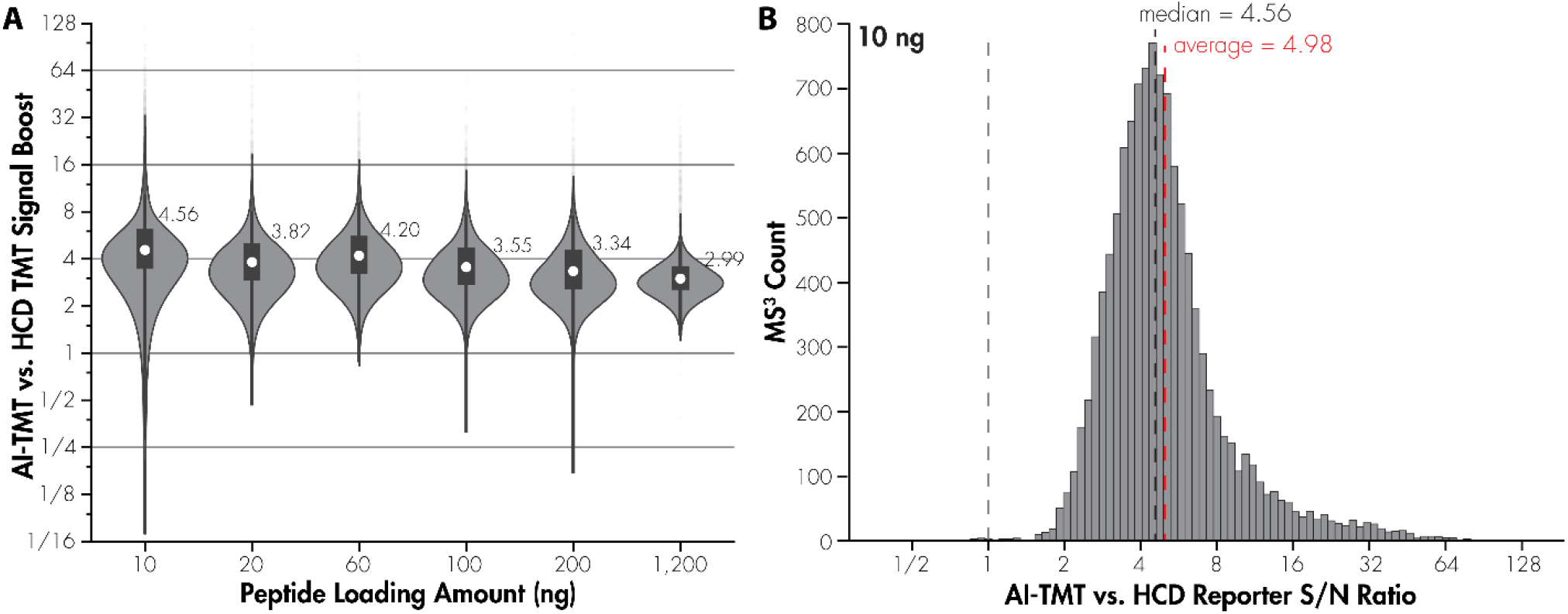
Increasing the quantitative reporter ion intensities for low-loading peptide injections. **A**, For a range of TMT-labeled peptide loading amounts, from 1,200 ng down to 10 ng, the ratio of reporter ion intensities with AI-TMT vs. HCD is shown. **B**, For the 10 ng peptide loading amount experiment, the distribution of AI-TMT vs. HCD reporter ion intensities is shown.

**Figure 3.**
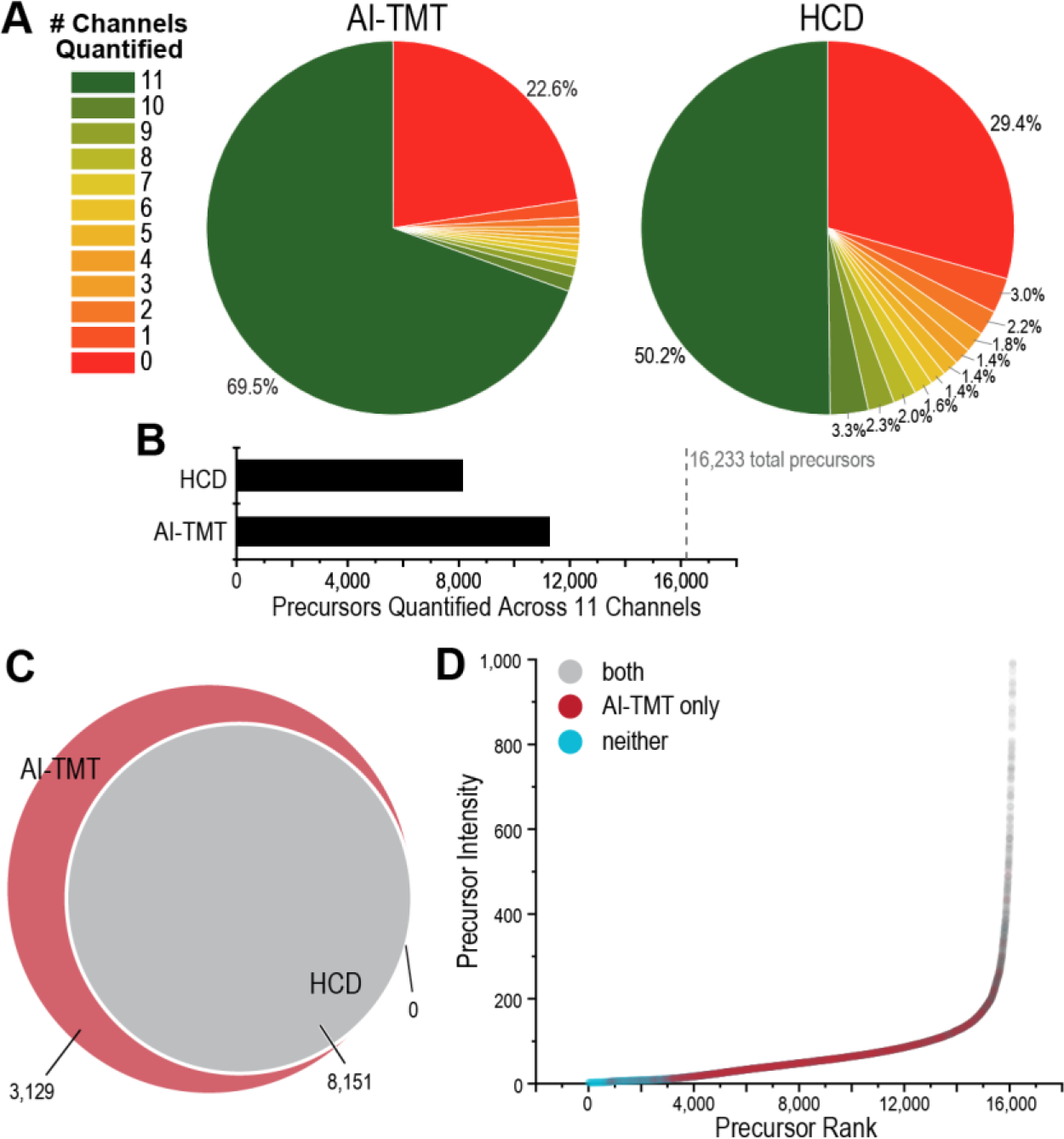
AI-TMT increases the dynamic range of peptide quantification. **A**, For a 10 ng peptide loading amount experiment, pie charts of the number of reporter ions quantified for AI-TMT and HCD SPS MS^3^ are shown (out of 11 total channels). **B**, Precursors fully quantified across all 11 channels. **C**, Venn diagram of precursors fully quantified across all 11 channels. **D**, The precursors fully quantified by AI-TMT (red) or both AI-TMT and HCD (grey) are shown, as well as the precursors that were not fully quantified (blue), ranked by their MS^1^ level precursor ion intensity.

We wanted to ensure that peptide quantification between reporter channels, one primary benefit of multiplexing with TMTs, is accurate and precise when using our AI-TMT platform. To interrogate this, we performed back-to-back experiments on precursors within the TMT 11-plex yeast triple knockout (TKO) sample. In back-to-back scans, we sampled the same precursor ion for collisional activation and isolated the same ten fragment ions for SPS-MS^3^ quantification, subjecting the ions to either HCD or AI-TMT activation. AI-TMT enables quantification of more peptide precursors regardless of what S/N threshold is applied (**Figure 4A**). Plotting the deviation between a peptide’s TMT channel ratios (i.e. 127c/126) from the mean ratio between those channels illustrates how AI-TMT enables more precise quantitation of peptides. The steepness of the curves in Figure 4B in combination with the number of MS^3^ scans above a given S/N cut-off help illustrate the precision of AI-TMT relative to HCD SPS-MS^3^.

**Figure 4.**
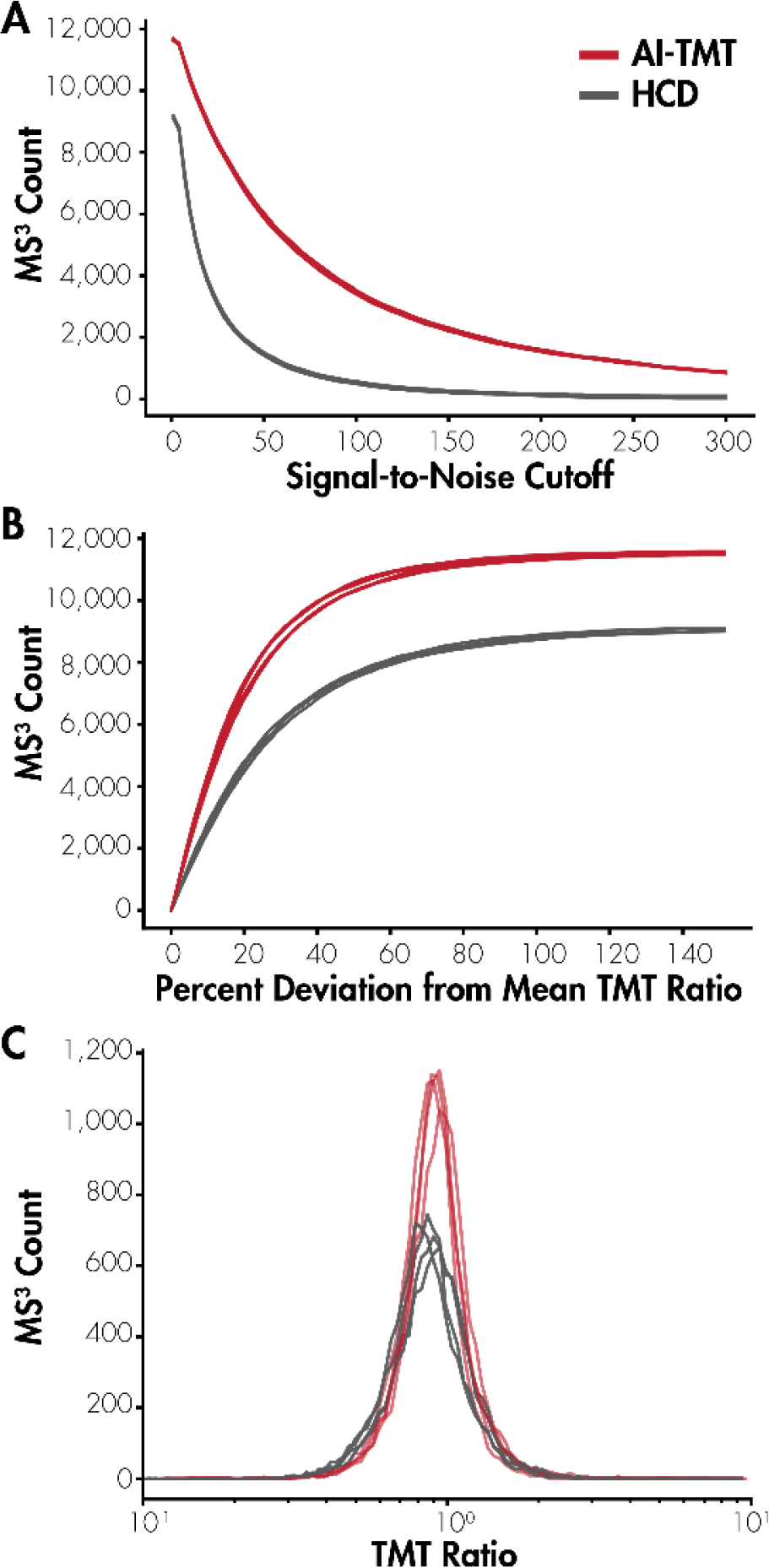
Quantitative accuracy and signal-to-noise distributions for peptides with AI-TMT and HCD SPS MS^3^. **A**, The distribution of the number of MS^3^ scans across the total TMT reporter ion signal-to-noise, summed across all channels. **B**, The distribution of the number of MS^3^ scans across the percent deviation from the mean TMT ratio. **C**, The number of quantifiable MS^3^ scans in both the intended channel and the TMT-126 channel and the associated TMT ratio between channels. Each line represents a different TMT channel, or ratio of TMT channels as with panel **4C**.

**Figure 4C** reveals the ratios of several TMT channels’ signal-to-noise to the TMT-126 channel. As all peptides are expected to carry equal abundance across channels (excluding the knockouts), we expect a ratio of 1, portrayed as 10^0^ in **Figure 4C** on a log_10_ scale. AI-TMT matches the quantitative accuracy as HCD while quantifying many more peptides (**Figure 4C**). As this experiment features back-to-back MS^3^ scans of the same precursor and the same 10 product ions activated with either HCD or AI-TMT, the increase in peptides quantified via AI-TMT must stem from zero values or peptides with values below the threshold of quantitation. Since our criteria for quantification of a TMT channel is S/N above zero, HCD is not resulting in Orbitrap MS^3^ detection of sufficient TMT reporter ions (∼5-10 ions) to yield a non-zero, quantitative value. This explains some of the non-linear benefits between peptide loading amount and relative boost of AI-TMT signal vs. HCD. Expanding these back-to-back yeast triple knockout experiments to single-cell levels of peptide injected (∼250 pg), we see comparable results. AI-TMT enables us to use 4-to 5-fold lower injection times than the conventional HCD approach while giving comparable TMT ion quantitative distributions across all peptides (**Figure 5** and **Figure S1**).

**Figure 5.**
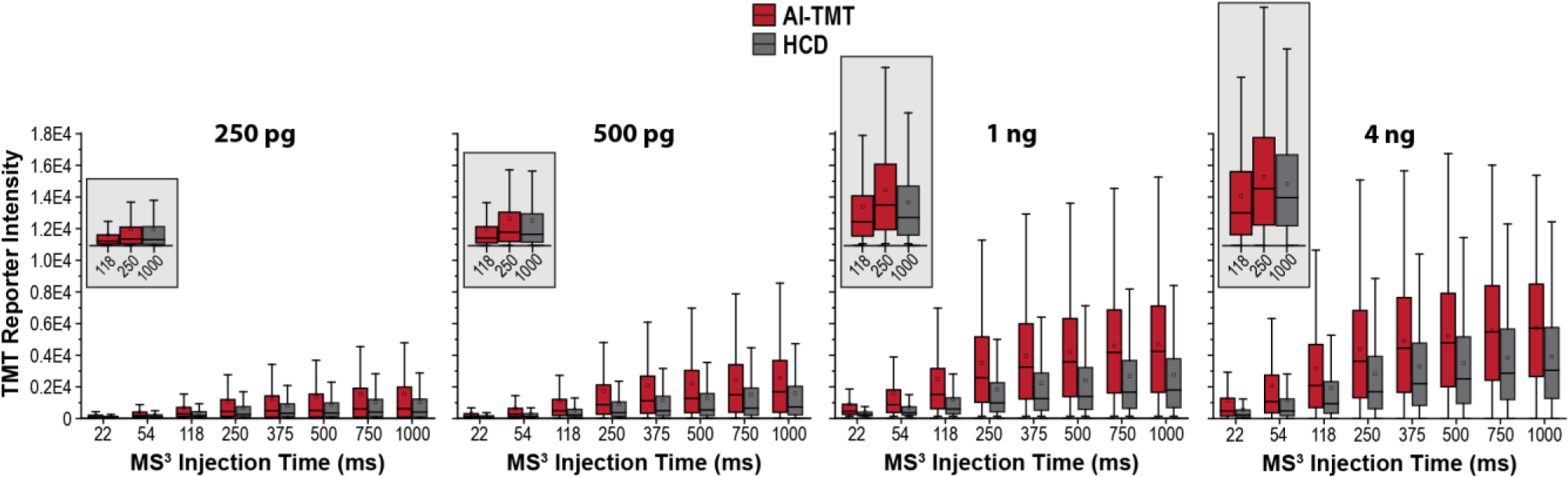
AI-TMT performance relative to HCD down to single-cell levels. Back-to-back experiments with either AI-TMT or HCD SPS MS^3^ illustrate the benefits of AI-TMT at varied peptide loading amounts and when decreasing the MS^3^ maximum ion injection time. Insets demonstrate that TMT reporter intensities using HCD at 1000 ms maximum injection time is comparable to that of AI-TMT at 118 ms and 250 ms.

### AI-TMT for single-cell proteomics

To further illustrate benefits of improved reporter ion yield, experiments of TMTpro 18-plex tagged HeLa and A549 single cells, prepared via nanoPOTS, were quantified at the MS^3^-level with HCD or IR photoactivation. Typically, single-cell proteomics researchers use exceptionally long MS^3^ injection times, often up to 750 ms or even longer, to ensure as much reporter ion signal as possible is detected. We notice that at only 118 ms injection time, AI-TMT can quantify a majority of the detected proteome; additional injection time does not help quantitation appreciably. Conversely, limiting MS^3^ injection time prevents quantification across most channels when performing activation by HCD (**Figure 6**). This inefficiency of HCD activation can be overcome by extending ion injection times to 750 ms. However, we note that similar performance is obtained with AI-TMT at 118 ms, roughly 4- to 5-fold less injection time. By using very long MS^3^ ion injection times, HCD can perform well, but it comes at a high cost to instrument duty cycle.

**Figure 6.**
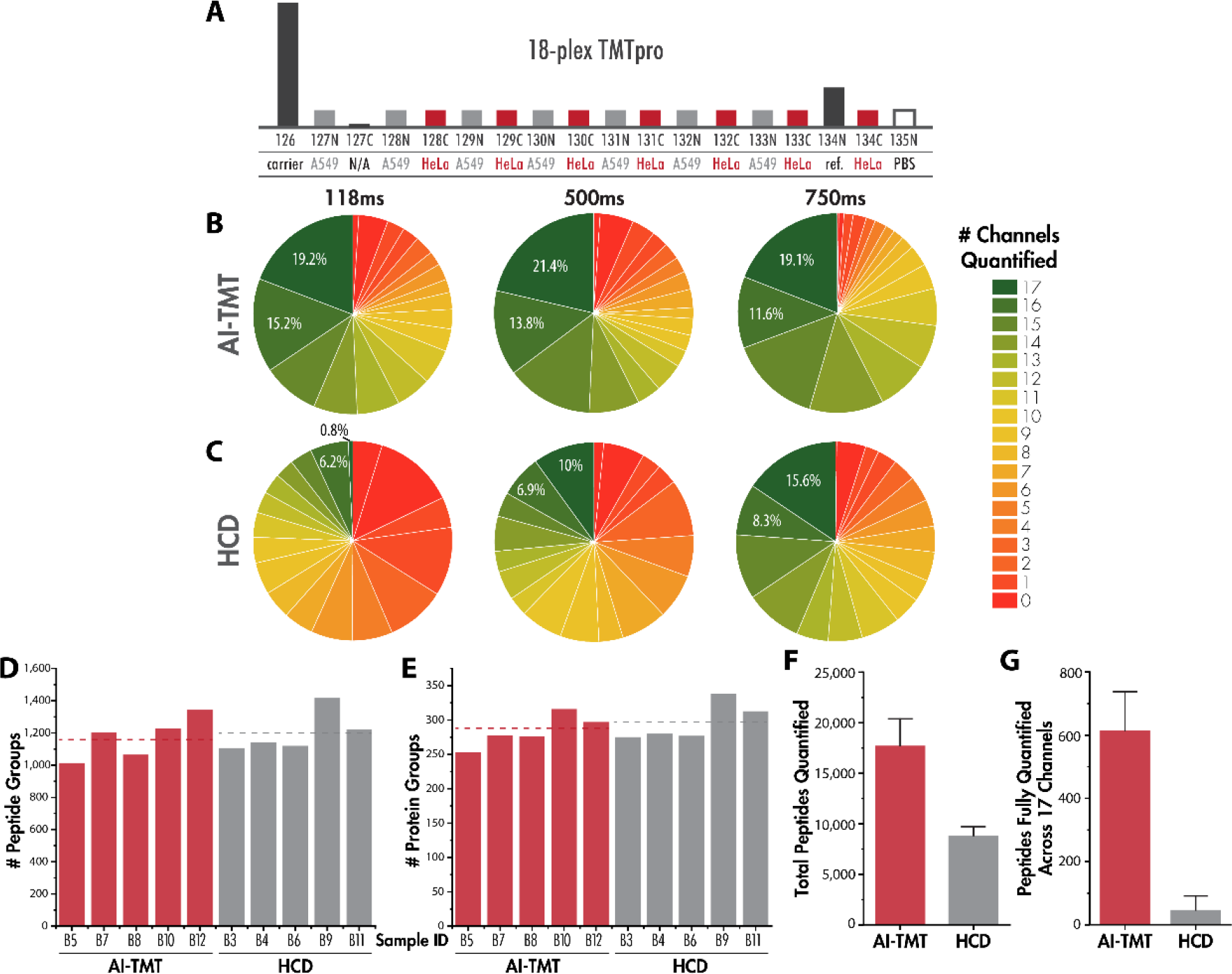
Improving the duty cycle of single-cell proteomics experiments. **A**, Depiction of the TMTpro 18plex channel configuration. Note that each sample contained the same number of carrier, PBS, reference, HeLa, and A549 channels, but in a different order. Pie charts of the number of peptides quantified across X channels for **B** AI-TMT and **C** HCD. The number of **D**, peptide groups and **E**, protein groups identified across five replicate experiments demonstrate comparable duty cycles. **F**, The total number of peptides quantified and the **G**, number of peptides quantified across all 17 channels are shown for these biological replicates.

In these comparative experiments, precursor peptides and their product ions are sampled multiple times per elution peak, activated by either HCD or AI-TMT in back-to-back MS^3^ scans. IR photoactivation identifies the same number of peptides and proteins as HCD, indicating no significant differences in scan speed exist between methods. Notably, AI-TMT quantifies about twice as many peptides overall as HCD and quantifies nearly 10-fold as many peptides across all 17 channels as HCD (**Figure 6**).

Given that AI-TMT can quantify 57% of peptides across all 17 TMT channels when given only 118 ms maximum ion injection times, and HCD MS^3^ quantifies less than 5% of peptides across all channels in a similar timeframe, we explored related MS^3^ data collection strategies. In duplicate LC-MS/MS experiments, we performed SPS-MS^3^ quantitation when MS^2^-level activation was performed with beam-type HCD and fragment ions were detected in the Orbitrap as well as when MS^2^-level activation was performed with resonant-type CAD and fragment ions were detected in the linear ion trap. For both experimental matrices, AI-TMT and HCD MS^3^ quantification were performed (**Figure 7**). Notably, AI-TMT quantifies 55% of peptides across all TMT channels in either scenario, whereas HCD quantifies only 25% of peptides across all TMT channels in either scenario. The MS^2^-level action and mass analyzer used in peptide identification do not seem to hold significant value for these single-cell proteomics results. Some researchers have posited that resonant-type CAD activation should be used for the MS^2^ activation stage in SPS-MS^3^ experiments as it induces peptide dissociation along fewer channels relative to HCD, yielding fewer but more abundant fragment ions.

**Figure 7.**
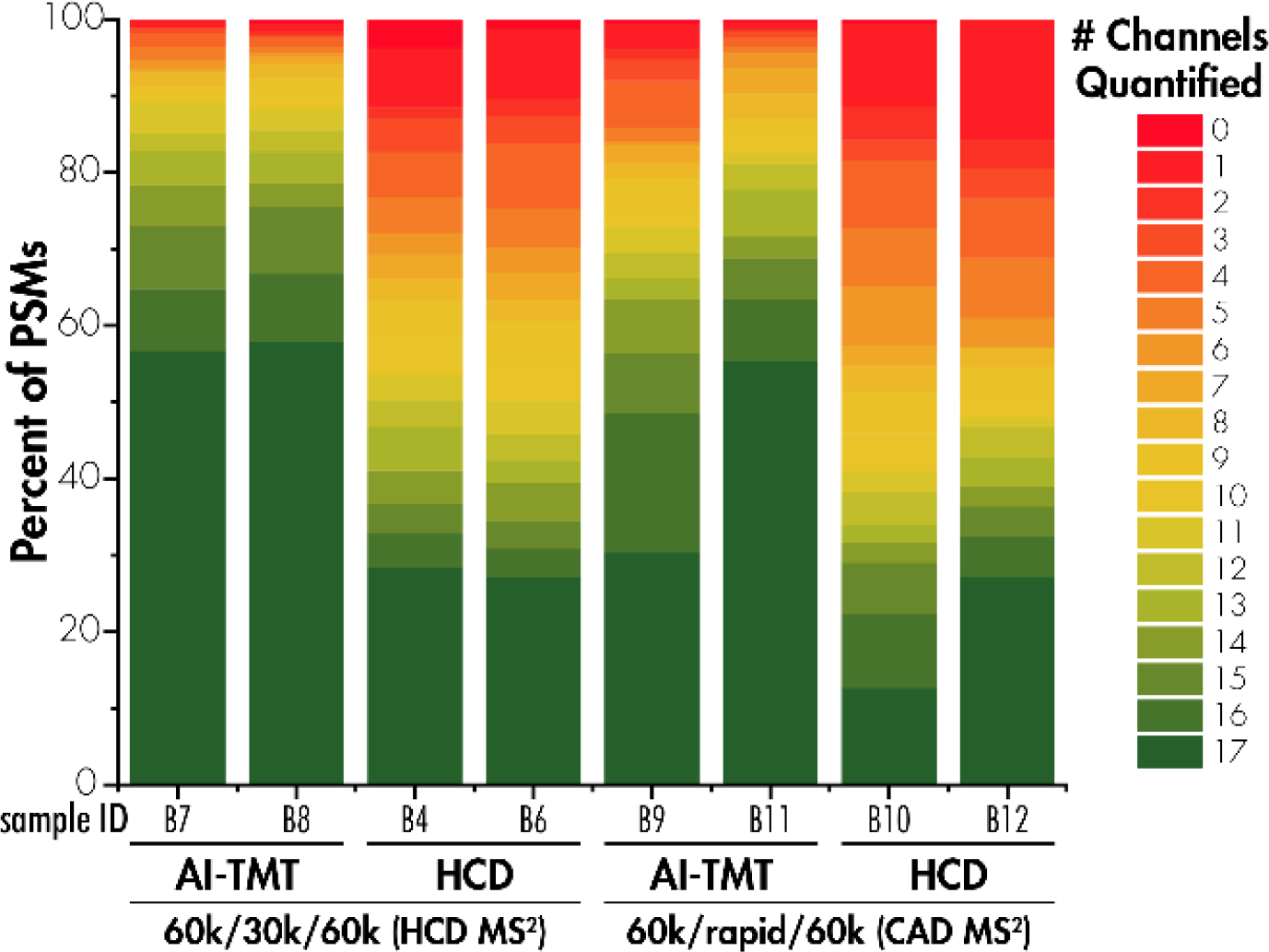
Comparison of single-cell proteomics data with alternative activation and mass analyzer strategies. TMTpro-18plex proteomic mixtures were fragmented with either CAD or HCD and ions were detected in either the ion trap (rapid) or Orbitrap (30,000 RP), respectively. AI-TMT and HCD SPS MS^3^ experiments were performed for both data collection strategies in duplicate with 250 ms max. ion injection times. Color of the bars represents the proportion of peptide spectral matches quantified across N number of channels (of 17 total channels).

## CONCLUSION

This advance not only improves the accuracy of single-cell protein quantification, but also enables the detection and quantification of proteins present at lower abundance, ultimately enriching the depth and precision of single-cell proteomic analyses.

Moving forward, there are several recent advances in MS instrumentation that AI-TMT complements and should lead to greater biological outcomes. In the past two years, real-time searching (RTS) has featured prominently in pushing the throughput of TMT-based proteomics.^10,36^ The scan acquisition regime of SPS-MS^3^ methods is inherently slow relative to MS/MS-only strategies, and Schweppe et al. addressed this slower duty cycle with Orbiter RTS-MS.^36,37^ A real-time database search is performed on every MS/MS scan and a quantitative MS^3^ is only triggered upon confident peptide identification, yielding methods with twice as efficient data acquisition. Real-time searching is only available on Orbitrap Eclipse and Orbitrap Ascend Tribrid platforms and our goal is to extend AI-TMT to these platforms.^38–40^ One extension of RTS is targeted pathway proteomics with GoDig, which allows up to 2000 ms ion injection times for SPS-MS^3^ quantitation of low abundance peptides.^19^ Incorporation of AI-TMT into the GoDig workflow may enable sampling of even more peptides for targeted, multiplexed quantitation.^19,41^ Schoof *et al*. also recently described a new method, RETICLE, which boasts benefits over MS^2^-based quantitation for single-cell proteomics applications.^38^ Precursors are sampled for a rapid, data-dependent ion-trap MS^2^ scan and a high-resolution Orbitrap MS^2^ scan is triggered upon RTS peptide identification. RETICLE outperforms MS^2^ quantitation through deeper proteome coverage.^38^ It will be interesting to see how AI-TMT performs for peptide quantitation at the MS^2^-level. We have shown previously that the benefits of AI-TMT over HCD hold at the MS^2^-level and this improvement should be magnified at the single-cell proteome level.^23^

Researchers have been poignant to comment that care must be taken in how much material is used in carrier channels as to not distort reporter ion statistics.^2,42–45^ By using AI-TMT, researchers can limit the amount of material in each channel and avoid carrier channel effects, as sensitivity is boosted 4- to 5-fold with AI-TMT. We intend to extend this platform toward biologically-relevant applications, for example within the heterogenous nature of single cancer cells, where extending the dynamic range of quantification to low-abundance proteins will prove useful.

This approach should also improve iTRAQ, DiLeu, and other labeling strategies that seek to increase the multiplexing capacity of single-cell proteomics and post-translational modification analysis.^22,46–50^ As TMT workflows advance to higher plex configurations, the initial precursor ion population will be split into many more channels and detection of reporter ions above the limit of quantitation for every channel will be even more challenging; AI-TMT is well poised to address this challenge.

## Supporting information

Supplemental Methods and Results

## ASSOCIATED CONTENT

### Supporting Information

Expanded materials and methods descriptions. **Figure S1**, method performance for a range of protein loading amounts; **Figure S2**, boosting peptide reporter ion quantitation at 500 pg loading amount; **Figure S3**, benefits of increased maximum ion injection time for AI-TMT and HCD experiments; **Figure S4**, AI-TMT enables quantification of low abundance peptide ions; **Figure S5**, samples used in this study; **Figure S6**, back-to-back analysis of using AI-TMT or HCD for generation of reporter ions from the TKO yeast standard; **Figure S7**, MS^3^ ion injection times during HCD and AI-TMT experiments.

## AUTHOR INFORMATION

### Author Contributions

Conceptualization, T.P.C., Y.R.L., R.T.K., and J.J.C.; Methodology, T.P.C., Y.R.L., R.T.K., and J.J.C.; Software, K.L.M., K.W.L., and T.P.C.; Validation, T.P.C., Y.R.L., K.L.M., M.S.W., J.D.H., G.C.M., J.E.P.S., R.T.K., and J.J.C.; Formal Analysis, T.P.C. and K.L.M.; Writing – Original Draft Preparation, T.P.C.; Writing – Review & Editing, T.P.C., Y.R.L., R.T.K., and J.J.C.; Visualization, T.P.C.; Supervision, J.J.C. and R.T.K.; Funding Acquisition, J.J.C. and R.T.K.

### Notes

Raw data files are available online on Chorus, project ID #1852 (https://chorusproject.org/pages/dashboard.html#/projects/all/1852/experiments).

## ACKNOWLEDGMENT

The authors thank other members of the Coon Lab for helpful discussions. This work was supported by the National Institute of General Medical Sciences of the National Institutes of Health (grant P41GM108538 to J.J.C.) and the National Human genome Research Institution through a training grant to the Genomic Science Training Program (grant T32HG002760 to T.M.P.-C.). T.M.P.-C. acknowledges the ACS Division of Analytical Chemistry and Agilent for support through a graduate fellowship.

## REFERENCES

(1) Kelly, R. T. Single-Cell Proteomics: Progress and Prospects. Mol. Cell. Proteomics 2020, 19 (11), 1739–1748. 10.1074/mcp.r120.002234.

(2) Ctortecka, C.; Mechtler, K. The Rise of Single-cell Proteomics. Analytical Science Advances 2021, 2 (3–4), 84–94. 10.1002/ANSA.202000152.

(3) Petrosius, V.; Schoof, E. M. Recent Advances in the Field of Single-Cell Proteomics. Transl Oncol 2023, 27 (October 2022), 101556. 10.1016/j.tranon.2022.101556.

(4) Peters-Clarke, T. M.; Coon, J. J.; Riley, N. M. Instrumentation at the Leading Edge of Proteomics. chemRxiv 2023. 10.26434/chemrxiv-2023-8l72m.

(5) Altelaar, A. F. M.; Heck, A. J. R. Trends in Ultrasensitive Proteomics. Curr. Opin. Chem. Biol. 2012, 16 (1–2), 206–213. 10.1016/j.cbpa.2011.12.011.

(6) Budnik, B.; Levy, E.; Harmange, G.; Slavov, N. SCoPE-MS: Mass Spectrometry of Single Mammalian Cells Quantifies Proteome Heterogeneity during Cell Differentiation 06 Biological Sciences 0601 Biochemistry and Cell Biology 06 Biological Sciences 0604 Genetics. Genome Biol 2018, 19 (1), 1–12.

(7) Specht, H.; Emmott, E.; Petelski, A. A.; Huffman, R. G.; Perlman, D. H.; Serra, M.; Kharchenko, P.; Koller, A.; Slavov, N. Single-Cell Proteomic and Transcriptomic Analysis of Macrophage Heterogeneity Using SCoPE2. Genome Biol 2021, 22 (1), 1–27. 10.1186/s13059-021-02267-5.

(8) Polychronidou, M.; Hou, J.; Babu, M. M.; Liberali, P.; Amit, I.; Deplancke, B.; Lahav, G.; Itzkovitz, S.; Mann, M.; Saez-Rodriguez, J.; Theis, F.; Eils, R. Single-Cell Biology: What Does the Future Hold? Mol Syst Biol 2023, e11799. 10.15252/MSB.202311799.

(9) Schoof, E. M.; Furtwängler, B.; Üresin, N.; Rapin, N.; Savickas, S.; Gentil, C.; Lechman, E.; Keller, U. auf dem; Dick, J. E.; Porse, B. T. Quantitative Single-Cell Proteomics as a Tool to Characterize Cellular Hierarchies. Nat Commun 2021, 12 (1), 1–15. 10.1038/s41467-021-23667-y.

(10) Furtwängler, B.; Üresin, N.; Motamedchaboki, K.; Huguet, R.; Lopez-Ferrer, D.; Zabrouskov, V.; Porse, B. T.; Schoof, E. M. Real-Time Search-Assisted Acquisition on a Tribrid Mass Spectrometer Improves Coverage in Multiplexed Single-Cell Proteomics. Molecular and Cellular Proteomics 2022, 21 (4), 100219. 10.1016/J.MCPRO.2022.100219/ATTACHMENT/84D9CAC4-0594-4038-92E1-9965B57C3B63/MMC5.PDF.

(11) Bouwmeester, R.; Gabriels, R.; Hulstaert, N.; Martens, L.; Degroeve, S. DeepLC Can Predict Retention Times for Peptides That Carry As-yet Unseen Modifications. Nat Methods 2021, 18 (11), 1363–1369. 10.1038/s41592-021-01301-5.

(12) Zhu, Y.; Piehowski, P. D.; Zhao, R.; Chen, J.; Shen, Y.; Moore, R. J.; Shukla, A. K.; Petyuk, V. A.; Campbell-Thompson, M.; Mathews, C. E.; Smith, R. D.; Qian, W. J.; Kelly, R. T. Nanodroplet Processing Platform for Deep and Quantitative Proteome Profiling of 10–100 Mammalian Cells. Nat Commun 2018, 9 (1), 1–10. 10.1038/s41467-018-03367-w.

(13) Matsumoto, C.; Shao, X.; Bogosavljevic, M.; Chen, L.; Gao, Y. Automated Container-Less Cell Processing Method for Single-Cell Proteomics. bioRxiv 2022, 12, 2022.07.26.501646. 10.1101/2022.07.26.501646.

(14) Li, J.; Van Vranken, J. G.; Pontano Vaites, L.; Schweppe, D. K.; Huttlin, E. L.; Etienne, C.; Nandhikonda, P.; Viner, R.; Robitaille, A. M.; Thompson, A. H.; Kuhn, K.; Pike, I.; Bomgarden, R. D.; Rogers, J. C.; Gygi, S. P.; Paulo, J. A. TMTpro Reagents: A Set of Isobaric Labeling Mass Tags Enables Simultaneous Proteome-Wide Measurements across 16 Samples. Nat Methods 2020. 10.1038/S41592-020-0781-4.

(15) Thompson, A.; Wölmer, N.; Koncarevic, S.; Selzer, S.; Böhm, G.; Legner, H.; Schmid, P.; Kienle, S.; Penning, P.; Höhle, C.; Berfelde, A.; Martinez-Pinna, R.; Farztdinov, V.; Jung, S.; Kuhn, K.; Pike, I. TMTpro: Design, Synthesis, and Initial Evaluation of a Proline-Based Isobaric 16-Plex Tandem Mass Tag Reagent Set. Anal Chem 2019, 91 (24), 15941–15950. 10.1021/ACS.ANALCHEM.9B04474.

(16) Li, J.; Cai, Z.; Bomgarden, R. D.; Pike, I.; Kuhn, K.; Rogers, J. C.; Roberts, T. M.; Gygi, S. P.; Paulo, J. A. TMTpro-18plex: The Expanded and Complete Set of TMTpro Reagents for Sample Multiplexing. J Proteome Res 2021, 20 (5), 2964–2972. 10.1021/acs.jproteome.1c00168.

(17) Zecha, J.; Satpathy, S.; Kanashova, T.; Avanessian, S. C.; Kane, M. H.; Clauser, K. R.; Mertins, P.; Carr, S. A.; Kuster, B. TMT Labeling for the Masses: A Robust and Cost-Efficient, in-Solution Labeling Approach. Molecular and Cellular Proteomics 2019, 18 (7), 1468–1478. 10.1074/MCP.TIR119.001385/ATTACHMENT/8DF2EBB4-9EFD-4D2F-823F-AC8A09C4A660/MMC1.ZIP.

(18) Shuken, S. R. An Introduction to Mass Spectrometry-Based Proteomics. J Proteome Res 2023. 10.1021/ACS.JPROTEOME.2C00838.

(19) Yu, Q.; Liu, X.; Keller, M. P.; Navarrete-Perea, J.; Zhang, T.; Fu, S.; Vaites, L. P.; Shuken, S. R.; Schmid, E.; Keele, G. R.; Li, J.; Huttlin, E. L.; Rashan, E. H.; Simcox, J.; Churchill, G. A.; Schweppe, D. K.; Attie, A. D.; Paulo, J. A.; Gygi, S. P. Sample Multiplexing-Based Targeted Pathway Proteomics with Real-Time Analytics Reveals the Impact of Genetic Variation on Protein Expression. Nat Commun 2023, 14 (1), 1–16. 10.1038/s41467-023-36269-7.

(20) Ting, L.; Rad, R.; Gygi, S. P.; Haas, W. MS3 Eliminates Ratio Distortion in Isobaric Multiplexed Quantitative Proteomics. Nat Methods 2011, 8 (11), 937–940. 10.1038/nmeth.1714.

(21) McAlister, G. C.; Nusinow, D. P.; Jedrychowski, M. P.; Wühr, M.; Huttlin, E. L.; Erickson, B. K.; Rad, R.; Haas, W.; Gygi, S. P. MultiNotch MS3 Enables Accurate, Sensitive, and Multiplexed Detection of Differential Expression across Cancer Cell Line Proteomes. Anal Chem 2014, 86 (14), 7150–7158. 10.1021/AC502040V.

(22) McAlister, G. C.; Huttlin, E. L.; Haas, W.; Ting, L.; Jedrychowski, M. P.; Rogers, J. C.; Kuhn, K.; Pike, I.; Grothe, R. A.; Blethrow, J. D.; Gygi, S. P. Increasing the Multiplexing Capacity of TMTs Using Reporter Ion Isotopologues with Isobaric Masses. Anal Chem 2012, 84 (17), 7469–7478. 10.1021/AC301572T/SUPPL_FILE/AC301572T_SI_001.PDF.

(23) Lee, K. W.; Peters-Clarke, T. M.; Mertz, K. L.; McAlister, G. C.; Syka, J. E. P.; Westphall, M. S.; Coon, J. J. Infrared Photoactivation Boosts Reporter Ion Yield in Isobaric Tagging. Anal Chem 2022, 94 (7), 3328–3334. 10.1021/ACS.ANALCHEM.1C05398/ASSET/IMAGES/MEDIUM/AC1C05398_M001.GIF.

(24) Liang, Y.; Truong, T.; Saxton, A. J.; Boekweg, H.; Payne, S. H.; Van Ry, P. M.; Kelly, R. T. HyperSCP: Combining Isotopic and Isobaric Labeling for Higher Throughput Single-Cell Proteomics. Anal Chem 2023, 95 (20), 8020–8027. 10.1021/acs.analchem.3c00906.

(25) Woo, J.; Williams, S. M.; Markillie, L. M.; Feng, S.; Tsai, C. F.; Aguilera-Vazquez, V.; Sontag, R. L.; Moore, R. J.; Hu, D.; Mehta, H. S.; Cantlon-Bruce, J.; Liu, T.; Adkins, J. N.; Smith, R. D.; Clair, G. C.; Pasa-Tolic, L.; Zhu, Y. High-Throughput and High-Efficiency Sample Preparation for Single-Cell Proteomics Using a Nested Nanowell Chip. Nat Commun 2021, 12 (1), 1–11. 10.1038/s41467-021-26514-2.

(26) Zhu, Y.; Podolak, J.; Zhao, R.; Shukla, A. K.; Moore, R. J.; Thomas, G. V.; Kelly, R. T. Proteome Profiling of 1 to 5 Spiked Circulating Tumor Cells Isolated from Whole Blood Using Immunodensity Enrichment, Laser Capture Microdissection, Nanodroplet Sample Processing, and Ultrasensitive NanoLC-MS. Anal Chem 2018, 90 (20), 11756–11759. 10.1021/acs.analchem.8b03268.

(27) Hailemariam, M.; Eguez, R. V.; Singh, H.; Bekele, S.; Ameni, G.; Pieper, R.; Yu, Y. S-Trap, an Ultrafast Sample-Preparation Approach for Shotgun Proteomics. J Proteome Res 2018, 17 (9), 2917–2924. 10.1021/acs.jproteome.8b00505.

(28) Peters-Clarke, T.; Schauer, K.; Riley, N.; Lodge, J.; Westphall, M.; Coon, J. Optical Fiber-Enabled Photoactivation of Peptides and Proteins. Anal Chem 2020, 92 (18), 12363–12370. 10.1021/acs.analchem.0c02087.

(29) Riley, N. M.; Westphall, M. S.; Hebert, A. S.; Coon, J. J. Implementation of Activated Ion Electron Transfer Dissociation on a Quadrupole-Orbitrap-Linear Ion Trap Hybrid Mass Spectrometer. Anal Chem 2017, 89, 6358–6366. 10.1021/acs.analchem.7b00213.

(30) Shishkova, E.; Hebert, A. S.; Westphall, M. S.; Coon, J. J. Ultra-High Pressure (>30,000 Psi) Packing of Capillary Columns Enhancing Depth of Shotgun Proteomic Analyses. Anal Chem 2018, 90 (19), 11503–11508. 10.1021/acs.analchem.8b02766.

(31) Brademan, D. R.; Riley, N. M.; Kwiecien, N. W.; Coon, J. J. Interactive Peptide Spectral Annotator: A Versatile Web-Based Tool for Proteomic Applications. Molecular and Cellular Proteomics 2019, 14, 193–201.

(32) Kovalchik, K. A.; Moggridge, S.; Chen, D. D. Y.; Morin, G. B.; Hughes, C. S. Parsing and Quantification of Raw Orbitrap Mass Spectrometer Data Using RawQuant. J Proteome Res 2018, 17 (6), 2237–2247. 10.1021/acs.jproteome.8b00072.

(33) Wenger, C. D.; Phanstiel, D. H.; Lee, M. V.; Bailey, D. J.; Coon, J. J. COMPASS: A Suite of Pre-and Post-Search Proteomics Software Tools for OMSSA. Proteomics 2011. 10.1002/pmic.201000616.

(34) Wenger, C. D.; Coon, J. J. A Proteomics Search Algorithm Specifically Designed for High-Resolution Tandem Mass Spectra. J Proteome Res 2013, 12 (3), 1377–1386. 10.1021/pr301024c.

(35) Bailey, D. J.; McDevitt, M. T.; Westphall, M. S.; Pagliarini, D. J.; Coon, J. J. Intelligent Data Acquisition Blends Targeted and Discovery Methods. 2014, 13 (4), 2152–2161.

(36) Schweppe, D. K.; Eng, J. K.; Yu, Q.; Bailey, D.; Rad, R.; Navarrete-Perea, J.; Huttlin, E. L.; Erickson, B. K.; Paulo, J. A.; Gygi, S. P. Full-Featured, Real-Time Database Searching Platform Enables Fast and Accurate Multiplexed Quantitative Proteomics. J Proteome Res 2020, 19 (5), 2026–2034.

(37) McAlister, G. C.; Nusinow, D. P.; Jedrychowski, M. P.; Wühr, M.; Huttlin, E. L.; Erickson, B. K.; Rad, R.; Haas, W.; Gygi, S. P. MultiNotch MS3 Enables Accurate, Sensitive, and Multiplexed Detection of Differential Expression across Cancer Cell Line Proteomes. Anal Chem 2014, 86 (14), 7150–7158.

(38) Furtwängler, B.; Üresin, N.; Motamedchaboki, K.; Huguet, R.; Lopez-Ferrer, D.; Zabrouskov, V.; Porse, B. T.; Schoof, E. M. Real-Time Search-Assisted Acquisition on a Tribrid Mass Spectrometer Improves Coverage in Multiplexed Single-Cell Proteomics. Molecular and Cellular Proteomics 2022, 21 (4), 100219. 10.1016/J.MCPRO.2022.100219/ATTACHMENT/84D9CAC4-0594-4038-92E1-9965B57C3B63/MMC5.PDF.

(39) He, Y.; Shishkova, E.; Peters-Clarke, T. M.; Brademan, D. R.; Westphall, M. S.; Bergen, D.; Huang, J.; Huguet, R.; Senko, M. W.; Zabrouskov, V.; McAlister, G. C.; Coon, J. J. Evaluation of the Orbitrap Ascend Tribrid Mass Spectrometer for Shotgun Proteomics. Anal Chem 2023, 95 (28). 10.1021/acs.analchem.3c01155.

(40) Shuken, S. R.; McAlister, G. C.; Barshop, W. D.; Canterbury, J. D.; Bergen, D.; Huang, J.; Huguet, R.; Paulo, J. A.; Zabrouskov, V.; Gygi, S. P.; Yu, Q. Deep Proteomic Compound Profiling with the Orbitrap Ascend Tribrid Mass Spectrometer Using Tandem Mass Tags and Real-Time Search. Anal Chem 2023, 95 (41), 15180–15188. 10.1021/acs.analchem.3c01701.

(41) Yu, Q.; Xiao, H.; Jedrychowski, M. P.; Schweppe, D. K.; Navarrete-Perea, J.; Knott, J.; Rogers, J.; Chouchani, E. T.; Gygi, S. P. Sample Multiplexing for Targeted Pathway Proteomics in Aging Mice. Proc Natl Acad Sci U S A 2020, 117 (18), 9723–9732. 10.1073/PNAS.1919410117/SUPPL_FILE/PNAS.1919410117.SD05.XLSX.

(42) Cheung, T. K.; Lee, C. Y.; Bayer, F. P.; McCoy, A.; Kuster, B.; Rose, C. M. Defining the Carrier Proteome Limit for Single-Cell Proteomics. Nat Methods 2020, 18 (1), 76–83. 10.1038/s41592-020-01002-5.

(43) Specht, H.; Slavov, N. Optimizing Accuracy and Depth of Protein Quantification in Experiments Using Isobaric Carriers. J. Proteome Res. 2021, 20 (1), 880–887. 10.1021/acs.jproteome.0c00675.

(44) Ye, Z.; Batth, T. S.; Rüther, P.; Olsen, J. V. A Deeper Look at Carrier Proteome Effects for Single-Cell Proteomics. Commun Biol 2022, 5 (1), 1–8. 10.1038/s42003-022-03095-4.

(45) Ctortecka, C.; Stejskal, K.; Krššáková, G.; Mendjan, S.; Mechtler, K. Quantitative Accuracy and Precision in Multiplexed Single-Cell Proteomics. Anal Chem 2022, 94 (5), 2434–2443. 10.1021/acs.analchem.1c04174.

(46) Xiang, F.; Ye, H.; Chen, R.; Fu, Q.; Li, N. N,N-Dimethyl Leucines as Novel Isobaric Tandem Mass Tags for Quantitative Proteomics and Peptidomics. Anal Chem 2010, 82 (7), 2817–2825. 10.1021/ac902778d.

(47) Wang, D.; Ma, M.; Huang, J.; Gu, T.-J.; Cui, Y.; Li, M.; Wang, Z.; Zetterberg, H.; Li, L. Boost-DiLeu: Enhanced Isobaric N,N-Dimethyl Leucine Tagging Strategy for a Comprehensive Quantitative Glycoproteomic Analysis. Anal Chem 2022, 94 (34), 11773–11782. 10.1021/ACS.ANALCHEM.2C01773.

(48) P. Ma T.; Izrael-Tomasevic, A.; Mroue, R.; Budayeva, H.; Malhotra, S.; Raisner, R.; Evangelista, M.; M. Rose C.; S. Kirkpatrick D.; Yu, K. AzidoTMT Enables Direct Enrichment and Highly Multiplexed Quantitation of Proteome-Wide Functional Residues. J Proteome Res 2023, 22 (7), 2218–2231. 10.1021/acs.jproteome.2c00703.

(49) Frost, D. C.; Feng, Y.; Li, L. 21-Plex DiLeu Isobaric Tags for High-Throughput Quantitative Proteomics. Anal Chem 2020, 92 (12), 8228–8234. 10.1021/acs.analchem.0c00473.

(50) Derks, J.; Leduc, A.; Wallmann, G.; Huffman, R. G.; Willetts, M.; Khan, S.; Specht, H.; Ralser, M.; Demichev, V.; Slavov, N. Increasing the Throughput of Sensitive Proteomics by PlexDIA. Nat Biotechnol 2022, 41, 1–10. 10.1038/s41587-022-01389-w.

