## Supplemental Methods and Results for "Boosting the Sensitivity of Quantitative Single-Cell Proteomics with Activated lon-Tandem Mass Tags (AI-TMT)"

**Figure S1.** Method performance for a range of peptide loading amounts. ........ S2

**Figure S2.** Boosting peptide reporter ion quantitation at 500 pg loading amount ........ S3

**Figure S3.** Benefits of increased maximum ion injection time for AI-TMT (top) and HCD (bottom) experiments ........ S4

**Figure S4.** AI-TMT enables quantification of low abundance peptide ions ........ S5

**Figure S5.** Samples used in this study ........ S6

**Figure S6.** Back-to-back analysis of using AI-TMT or HCD for generation of reporter ions from TKO yeast standard ........ S7

**Figure S7.** MS3 ion injection times during HCD and AI-TMT experiments ........ S8

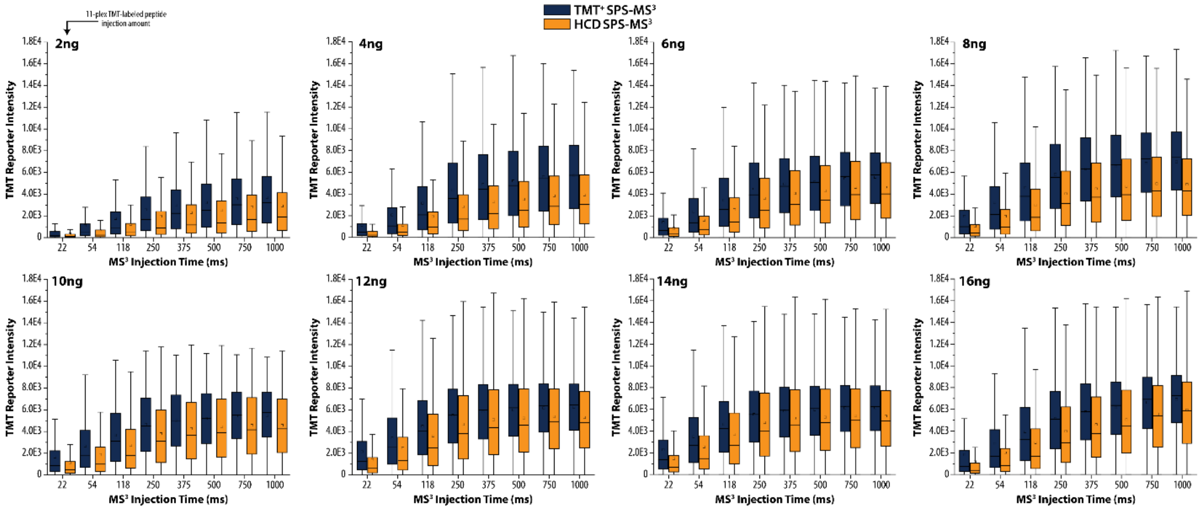

**Supp Fig. S1. Method performance for a range of peptide loading amounts.** Back-to-back experiments with either AI-TMT or HCD SPS-MS$^3$ illustrate the benefits of AI-TMT at varied peptide loading amounts, from 2 ng to 16 ng, and when decreasing the MS$^3$ maximum ion injection time, from 1000 ms down to 22 ms. AI-TMT is shown in dark blue and HCD is shown in light orange.

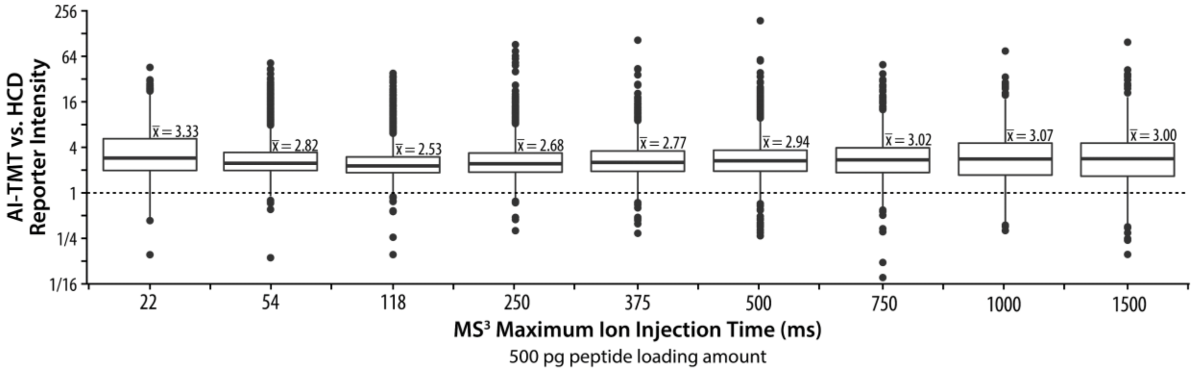

**Supp Fig. S2. Boosting peptide reporter ion quantitation at 500 pg loading amount.** The ratio of cumulative reporter ion intensities generated with AI-TMT vs. HCD is shown for 500 pg peptide loading amounts at varied $MS^3$ maximum ion injection time, from 1500 ms down to 22 ms.

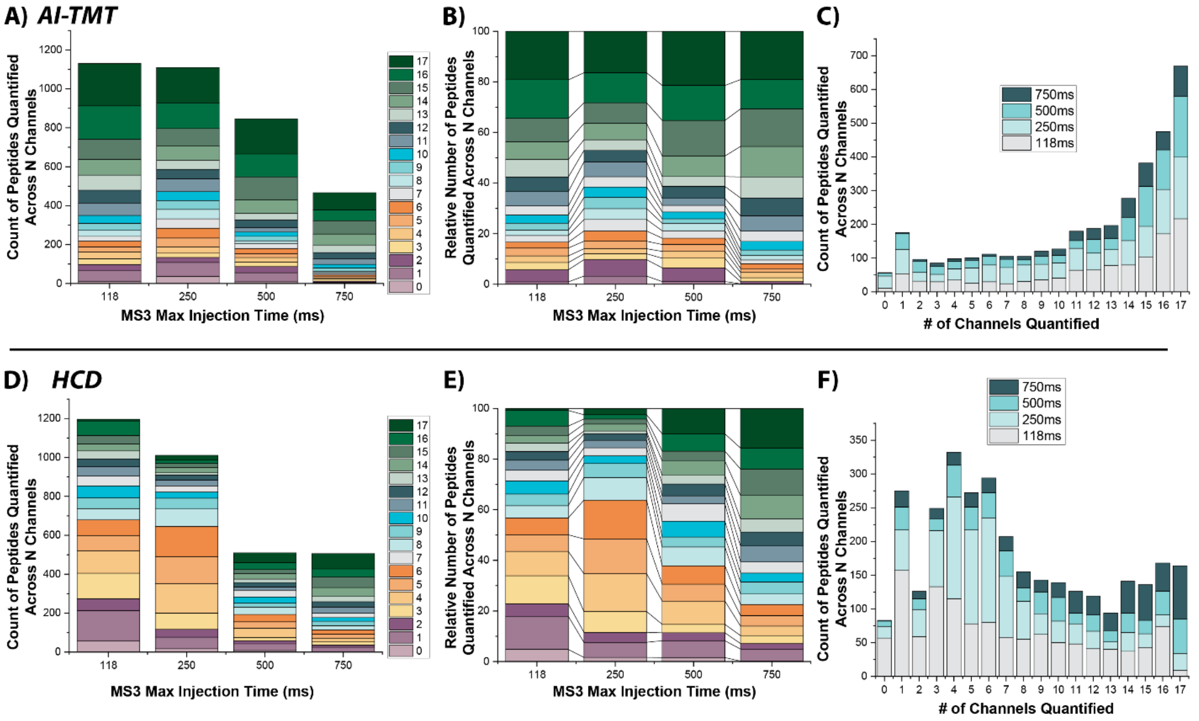

**Supp Fig. S3. Benefits of increased maximum ion injection time for AI-TMT (top) and HCD (bottom) experiments.** Increasing $MS^3$ ion injection time rescues HCD quantitation but is not necessary for AI-TMT.

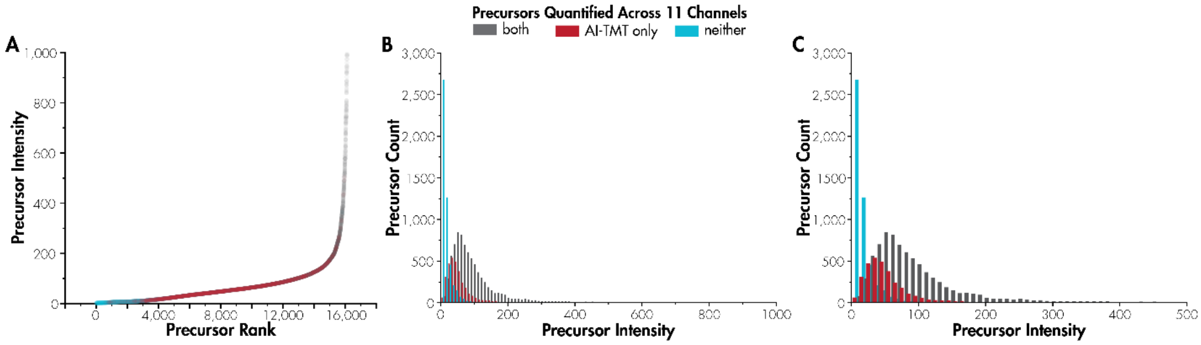

**Supp Fig. S4. AI-TMT enables quantification of low abundance peptide ions.** Back-to-back experiments with either AI-TMT or HCD SPS-MS$^3$ using 10 ng peptide loading illustrate the improvements of AI-TMT quantification over HCD. As an extension of Figure 3, this figure illustrates how low abundance peptides are more likely to be quantified across all eleven channels with AI-TMT.

| | 1 | 2 | 3 | 4 | 5 | 6 | 7 | 8 | 9 | 10 | 11 | 12 | 13 | 14 | 15 | 16 | 17 | 18 |
|---|---|---|---|---|---|---|---|---|---|---|---|---|---|---|---|---|---|---|
| **SampleWell**\Channel | **126** | **127N** | **127C** | **128N** | **128C** | **129N** | **129C** | **130N** | **130C** | **131N** | **131C** | **132N** | **132C** | **133N** | **133C** | **134N** | **134C** | **135N** |
| **Plate #1** | | | | | | | | | | | | | | | | | | |
| **C2** | Carrier | HeLa | NotUsed | PBS | HeLa | HeLa | HeLa | HeLa | HeLa | HeLa | HeLa | HeLa | HeLa | HeLa | HeLa | Reference | HeLa | HeLa |
| **C3** | Carrier | HeLa | NotUsed | HeLa | HeLa | HeLa | HeLa | HeLa | HeLa | HeLa | HeLa | HeLa | HeLa | HeLa | HeLa | Reference | HeLa | PBS |
| **C4** | Carrier | PBS | NotUsed | HeLa | HeLa | HeLa | HeLa | HeLa | HeLa | HeLa | HeLa | HeLa | HeLa | HeLa | HeLa | Reference | HeLa | HeLa |
| **C5** | Carrier | HeLa | NotUsed | HeLa | HeLa | HeLa | HeLa | PBS | HeLa | HeLa | HeLa | HeLa | HeLa | HeLa | HeLa | Reference | HeLa | HeLa |
| **C6** | Carrier | HeLa | NotUsed | HeLa | HeLa | HeLa | HeLa | HeLa | PBS | HeLa | HeLa | HeLa | HeLa | HeLa | HeLa | Reference | HeLa | HeLa |
| **C7** | Carrier | HeLa | NotUsed | HeLa | HeLa | HeLa | HeLa | HeLa | HeLa | HeLa | HeLa | PBS | HeLa | HeLa | HeLa | Reference | HeLa | HeLa |
| **C8** | Carrier | HeLa | NotUsed | HeLa | HeLa | HeLa | HeLa | HeLa | HeLa | PBS | HeLa | HeLa | HeLa | HeLa | HeLa | Reference | HeLa | HeLa |
| **C9** | Carrier | HeLa | NotUsed | HeLa | HeLa | HeLa | HeLa | HeLa | HeLa | HeLa | HeLa | HeLa | HeLa | HeLa | HeLa | Reference | HeLa | PBS |
| **C10** | Carrier | HeLa | NotUsed | HeLa | HeLa | HeLa | HeLa | HeLa | HeLa | HeLa | PBS | HeLa | HeLa | HeLa | HeLa | Reference | HeLa | HeLa |
| **C11** | Carrier | HeLa | NotUsed | HeLa | HeLa | PBS | HeLa | HeLa | HeLa | HeLa | HeLa | HeLa | HeLa | HeLa | HeLa | Reference | HeLa | HeLa |
| **C12** | Carrier | HeLa | NotUsed | HeLa | HeLa | HeLa | HeLa | HeLa | HeLa | HeLa | HeLa | HeLa | PBS | HeLa | HeLa | Reference | HeLa | HeLa |
| **C13** | Carrier | HeLa | NotUsed | HeLa | HeLa | HeLa | HeLa | HeLa | HeLa | HeLa | HeLa | PBS | HeLa | HeLa | HeLa | Reference | HeLa | HeLa |
| **C14** | Carrier | HeLa | NotUsed | HeLa | HeLa | HeLa | HeLa | HeLa | HeLa | PBS | HeLa | HeLa | HeLa | HeLa | HeLa | Reference | HeLa | HeLa |
| **C15** | Carrier | HeLa | NotUsed | HeLa | HeLa | HeLa | HeLa | HeLa | HeLa | HeLa | HeLa | PBS | HeLa | HeLa | HeLa | Reference | HeLa | HeLa |
| **C16** | Carrier | HeLa | NotUsed | HeLa | PBS | HeLa | HeLa | HeLa | HeLa | HeLa | HeLa | HeLa | HeLa | HeLa | HeLa | Reference | HeLa | HeLa |
| **C17** | Carrier | HeLa | NotUsed | HeLa | HeLa | HeLa | HeLa | HeLa | HeLa | HeLa | HeLa | HeLa | PBS | HeLa | HeLa | Reference | HeLa | HeLa |
| **Plate #2** | | | | | | | | | | | | | | | | | | |
| **B2** | Carrier | HeLa | NotUsed | HeLa | HeLa | A549 | A549 | HeLa | A549 | HeLa | HeLa | A549 | PBS | A549 | A549 | Reference | A549 | HeLa |
| **B3** | Carrier | A549 | NotUsed | HeLa | A549 | HeLa | A549 | PBS | A549 | HeLa | HeLa | A549 | HeLa | A549 | HeLa | Reference | A549 | HeLa |
| **B4** | Carrier | HeLa | NotUsed | HeLa | HeLa | A549 | A549 | HeLa | A549 | HeLa | HeLa | A549 | PBS | A549 | A549 | Reference | A549 | HeLa |
| **B5** | Carrier | HeLa | NotUsed | HeLa | HeLa | A549 | A549 | HeLa | A549 | A549 | A549 | PBS | HeLa | A549 | HeLa | Reference | HeLa | A549 |
| **B6** | Carrier | HeLa | NotUsed | HeLa | A549 | HeLa | A549 | HeLa | HeLa | PBS | A549 | A549 | HeLa | A549 | A549 | Reference | A549 | HeLa |
| **B7** | Carrier | HeLa | NotUsed | HeLa | A549 | HeLa | HeLa | A549 | A549 | HeLa | A549 | PBS | HeLa | A549 | A549 | Reference | A549 | HeLa |
| **B8** | Carrier | HeLa | NotUsed | HeLa | HeLa | A549 | A549 | HeLa | A549 | HeLa | HeLa | A549 | PBS | A549 | A549 | Reference | A549 | HeLa |
| **B9** | Carrier | A549 | NotUsed | HeLa | HeLa | A549 | HeLa | HeLa | A549 | HeLa | A549 | A549 | HeLa | PBS | HeLa | Reference | A549 | A549 |
| **B10** | Carrier | HeLa | NotUsed | PBS | HeLa | A549 | A549 | HeLa | A549 | HeLa | HeLa | HeLa | A549 | HeLa | A549 | Reference | A549 | A549 |
| **B11** | Carrier | A549 | NotUsed | A549 | HeLa | A549 | HeLa | HeLa | A549 | A549 | A549 | HeLa | HeLa | HeLa | A549 | Reference | HeLa | PBS |
| **B12** | Carrier | PBS | NotUsed | HeLa | A549 | HeLa | A549 | A549 | HeLa | A549 | HeLa | HeLa | HeLa | A549 | A549 | Reference | A549 | HeLa |
| **B13** | Carrier | A549 | NotUsed | HeLa | A549 | A549 | HeLa | HeLa | PBS | A549 | A549 | HeLa | HeLa | A549 | HeLa | Reference | A549 | HeLa |
| **B14** | Carrier | HeLa | NotUsed | A549 | HeLa | A549 | A549 | HeLa | HeLa | A549 | PBS | A549 | A549 | A549 | HeLa | Reference | HeLa | HeLa |
| **B15** | Carrier | A549 | NotUsed | HeLa | HeLa | HeLa | PBS | A549 | A549 | HeLa | A549 | HeLa | A549 | A549 | HeLa | Reference | A549 | HeLa |
| **B16** | Carrier | PBS | NotUsed | HeLa | A549 | HeLa | HeLa | A549 | HeLa | A549 | A549 | HeLa | A549 | HeLa | A549 | Reference | HeLa | A549 |
| **B17** | Carrier | HeLa | NotUsed | A549 | A549 | HeLa | A549 | HeLa | A549 | HeLa | A549 | HeLa | HeLa | A549 | HeLa | Reference | A549 | PBS |

**Supp Fig. S5. Samples used in this study.** Characterization of the cell type, control, reference channel, or carrier used in each multiplexed TMT experiment are shown.

TMT TKO Parking test: IRMPD 40% 20 V park vs. HCD 50nce; 10ng loading

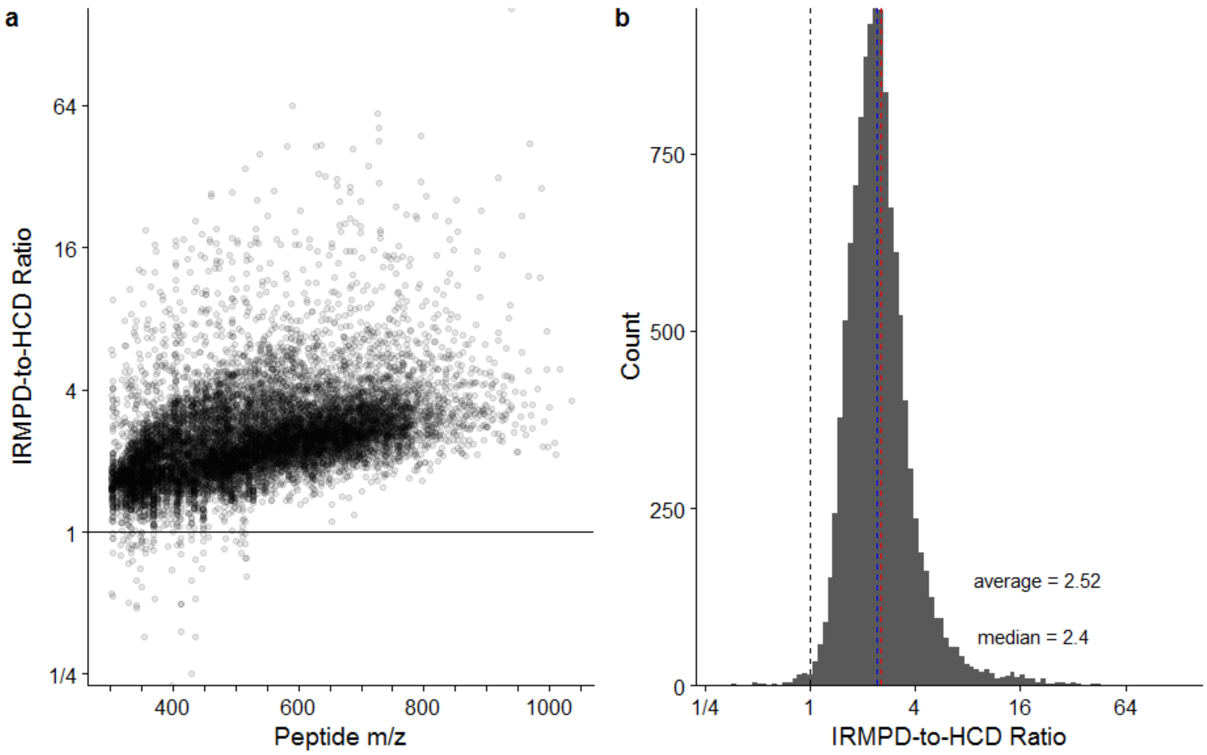

**Supp Fig. S6. Back-to-back analysis of using AI-TMT or HCD for generation of reporter ions from the TKO yeast standard.** Both experiments used HCD (30 nce) to generate fragments from individually isolated peptides and either 50 nce HCD or 40% IR for 5 ms to generate the TMT reporter from 10 synchronously selected peptide fragments. A 20 V parking waveform was applied during IRMPD to minimize loss of the reporter due to photodissociation. (a) Ratio of the reporter signal generated via AI-TMT over HCD vs yeast peptide *m/z*. (b) Histogram of reporter signal increase due to AI-TMT over HCD for all peptides.

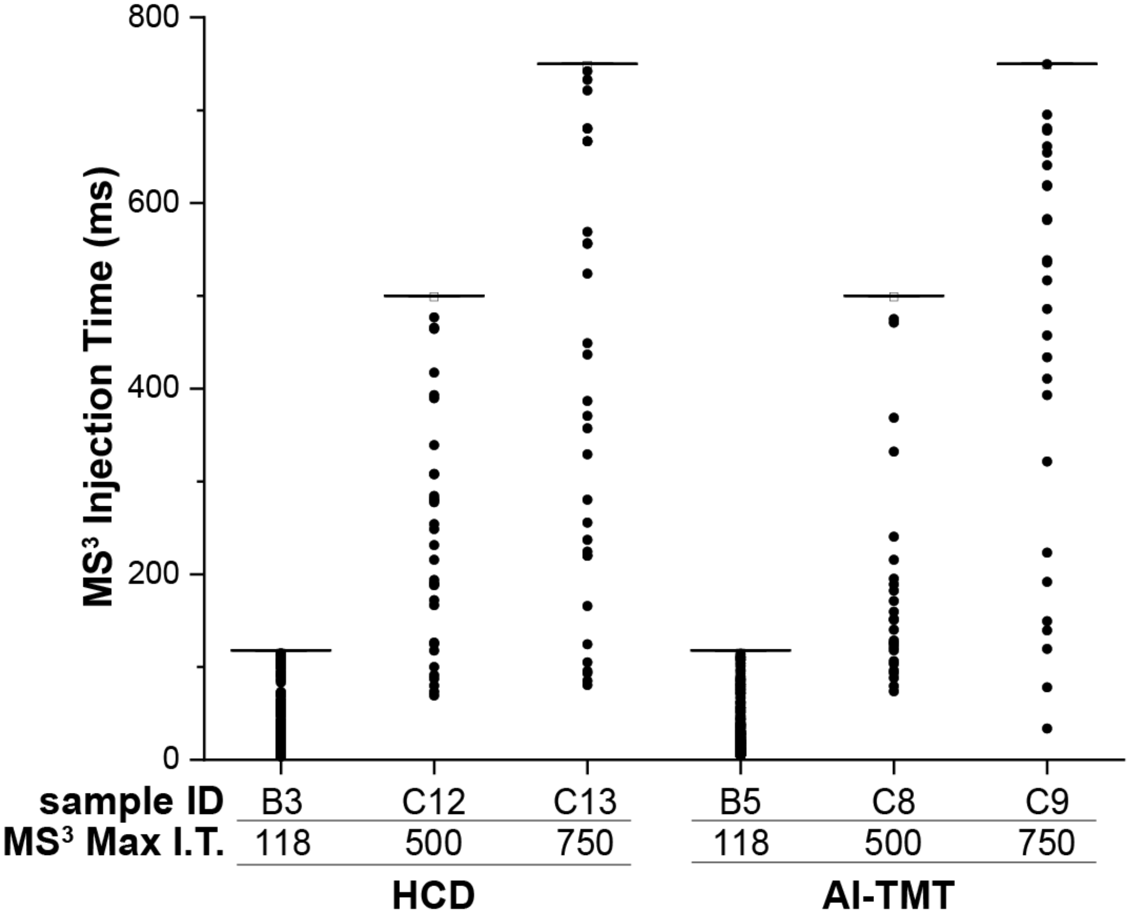

**Supp Fig. S7. MS$^3$ ion injection times during HCD and AI-TMT experiments.** In almost all cases, the maximum MS$^3$ injection time (MS$^3$ Max. I.T.) was reached.
